# Nanoscale Polarization of the Vaccinia Virus Entry Fusion Complex Drives Efficient Fusion

**DOI:** 10.1101/360073

**Authors:** Robert D. M. Gray, David Albrecht, Corina Beerli, Gary H. Cohen, Ricardo Henriques, Jason Mercer

**Author notes:** Equally contributing authors (Robert D. M. Gray, David Albrecht).

## Abstract

The spatial distribution of binding and fusion proteins on most viruses and the functional relevance of this organization remains largely unexplored. Employing super-resolution microscopy we define the nanoscale membrane architecture of the prototypic poxvirus, vaccinia. We show that binding and entry fusion complex (EFC) proteins are organized into distinct functional domains with fusion proteins polarized to the tips of virions. Repression of individual EFC components disrupted fusion protein polarization, correlating with a loss of fusion activity. Repression of vaccinia A27, a non-EFC protein implicated in fusion, revealed that disruption of EFC localization impacts virus fusion pore formation. We propose that the polarized distribution of EFCs is essential for poxvirus fusion efficiency.

## Introduction

For binding and fusion most viruses use between 1 and 5 proteins (1, 2). The *Poxviridae*, including variola, the causative agent of smallpox, monkeypox, and the smallpox vaccine, vaccinia virus, are an exception. All members of this virus family encode 4 proteins for binding and 11 proteins for fusion (3). The binding proteins (D8, H3, A26 and A27) are not individually essential (4–7). However, genetic repression of any of the 11 proteins required for fusion, collectively termed the entry fusion complex (EFC) (8, 9), results in formation of morphologically normal virions incompetent for hemifusion (A16, A21, F9, G3, G9, H2, J5 and 03) or full fusion (A28, L1 and L5) (8). All EFC components are transmembrane proteins and 9 (A16, A21, A28, G3, G9, H2, J5, L5 and O3) form a stable core complex with which L1 and F9 associate (8). For many viruses, fusion protein concentration is crucial for fusion efficiency (2, 10, 11). For human immunodeficiency virus (HIV-1), physical clustering of Env trimers during maturation correlates with entry competence (12). Although the spatial distribution of EFCs has not been explored, we noted an orientation bias in electron microscopy studies of poxvirus fusion. Virions preferentially bind the cell surface on their side, while fusion often appears to have occurred at virion tips, the functional significance of which has not been investigated (8, 13–15). Here, using a combination of super-resolution microscopy, single-particle analysis and a large collection of virus mutants, we investigate the spatial distribution of binding and fusion proteins on individual vaccinia virus (VACV) particles and how this correlates with fusion activity.

## Results

Combining structured illumination microscopy (SIM) (16) and single-particle averaging of hundreds of virions we have generated high-precision models of the average distribution of proteins in VACV mature virions MVs) (17, 18). We applied this imaging and analytical framework to investigate the localization of binding and EFC components on VACV MVs. For this, mCherry-tagged core protein A4 was used to identify virion position and orientation (Fig. 1A), and EGFP-tagged membrane protein A13 (19) as a viral membrane marker (Fig. 1A). VACV binding (A27 and D8) and EFC proteins (A21, A28, F9, J5, H2 and L1) were visualized by immunofluorescence. The localization models showed that MV binding proteins preferentially reside at the sides of virions, while all EFC components localize to the virion tips, independent of virion orientation (Fig. 1A, S1). Using these models, the polarity factor of each protein was quantified relative to the isotropically distributed core (A4) and membrane (A13) markers Fig. S2). Binding proteins were 1.2-fold enriched at the sides, and EFC proteins 1.7-fold enriched at the tips of virions (Fig. 1B).

The EFC is held together by a complex network of interactions (20). Repression of any core component results in disruption of the EFC into sub-complexes (21–23). Thus, we investigated whether the polarized distribution of binding and EFC components depends upon a fully intact EFC. Recombinant VACV MVs lacking EFC core components A28, G9 or O3 (24) were immunolabelled for D8 and L1, which respectively served as binding and fusion machinery markers. Localization models of D8 and L1 showed that D8 distribution was unaffected by loss of EFC components, while L1 was redistributed evenly around the MV membrane (Fig. 1C). Models of A21, A28, F9, H2 and J5 on A28(-), G9(-) and O3(-) MVs showed that all EFC components tested were depolarized in these mutants (Fig. S3A). No redistribution of D8 or EFC proteins was seen on virions lacking the binding protein H3 (Fig. 1C, Fig. S3A, B), indicating that H3 plays no role in MV membrane organization. Polarity factor quantification confirmed that deletion of EFC core components does not alter D8 localization, while EFC distribution is shifted from polarized to isotropic (Fig. 1D, Fig. S3B). As illustrated in Figure 1E, these results indicate that VACV binding and fusion machineries are organized as distinct functional domains within the viral membrane, and that the polarized distribution of the EFC relies on its intactness.

**Fig. 1.**
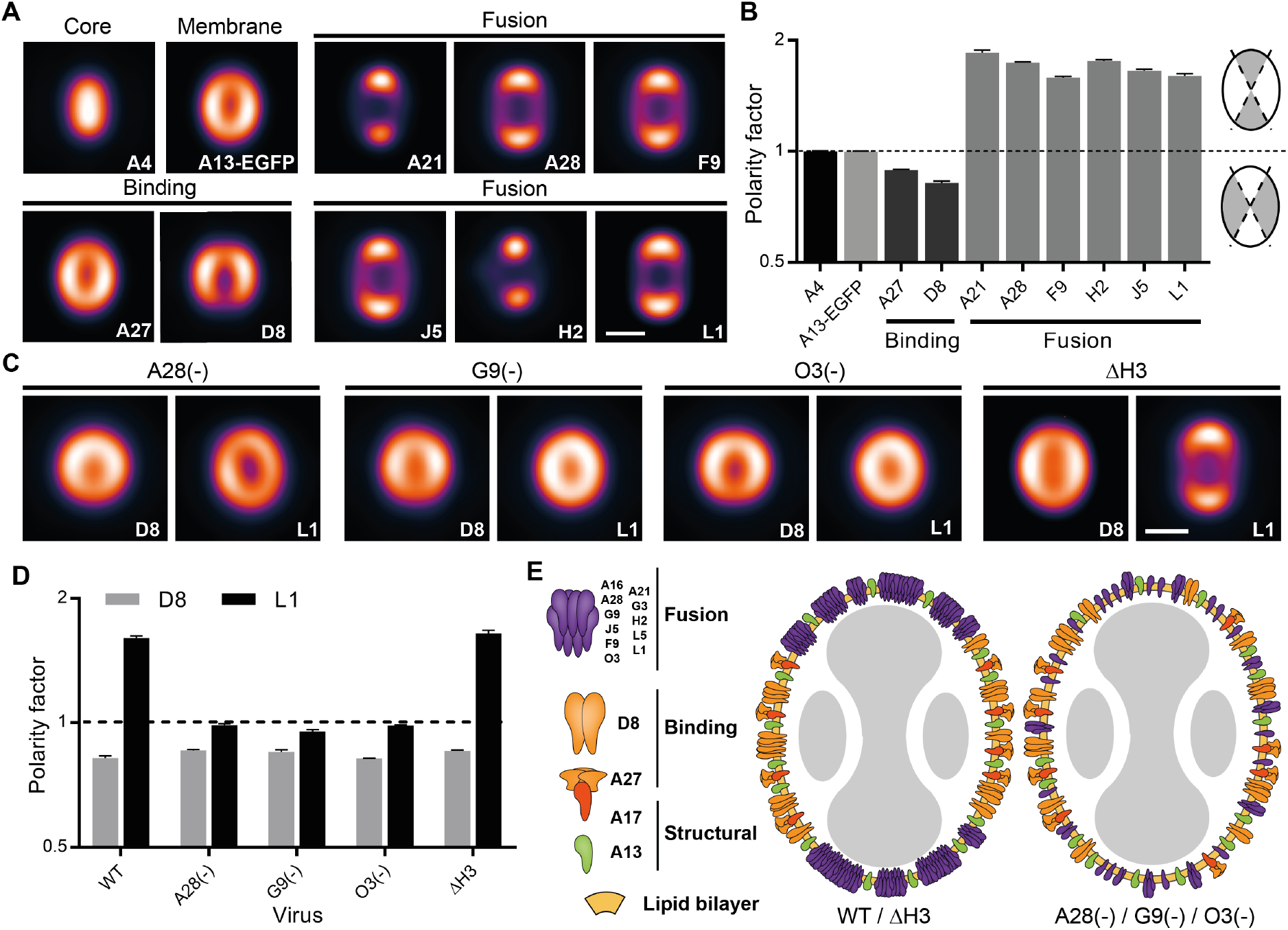
VACV binding and fusion proteins are organised into distinct domains. (A) Localization models of binding and fusion proteins on MVs. (B) Quantification of the polarity factor of proteins visualized in (A) on separately averaged subsets of 50 particles. A polarity factor > 1 corresponds to concentration at the tips of MVs; < 1 corresponds to concentration at the sides of MVs. (C) Localization models of D8 and L1 in EFC and binding mutant MVs. (D) Polarity factors for the models shown in (C) compared to WT for separately averaged subsets of 50 particles. (E) Illustration of VACV membrane protein organisation in WT/ΔH3 and fusion mutant MVs. Mean *±* SEM. Scale bars 200nm.

To extend our investigation of VACV binding and fusion machinery beyond population averaged localizations to high-resolution single virion analyses, we used direct stochastic optical reconstruction microscopy (dSTORM) (25). Immunolabeling of D8 and L1 was performed on WT, EFC mutant [A28(-), G9(-) and O3(-)], and binding protein mutant (ΔH3) MVs. Achieving resolutions in the order of tens of nanometers, dSTORM revealed that D8 and L1 were localized to distinct clusters in WT and ΔH3 MVs (Fig. 2A, B). Consistent with the localization models shown in Figure 1, D8 was distributed to the sides of MVs under all conditions, L1 was polarized to the tips of WT and ΔH3 MVs, and found throughout the membrane of A28(-), G9(-) and O3(-) virions (Fig. 2A, Fig. S4A). D8 clusters appeared unaffected in EFC mutant virions, while the clusters of L1 were largely disrupted (Fig. 2A). Similar patterns ofL1 and D8 distribution were observed on WT virions stained with fluorescently conjugated primary antibodies, ruling out antibody-induced clustering (Fig. S4B, C) (26).

**Fig. 2.**
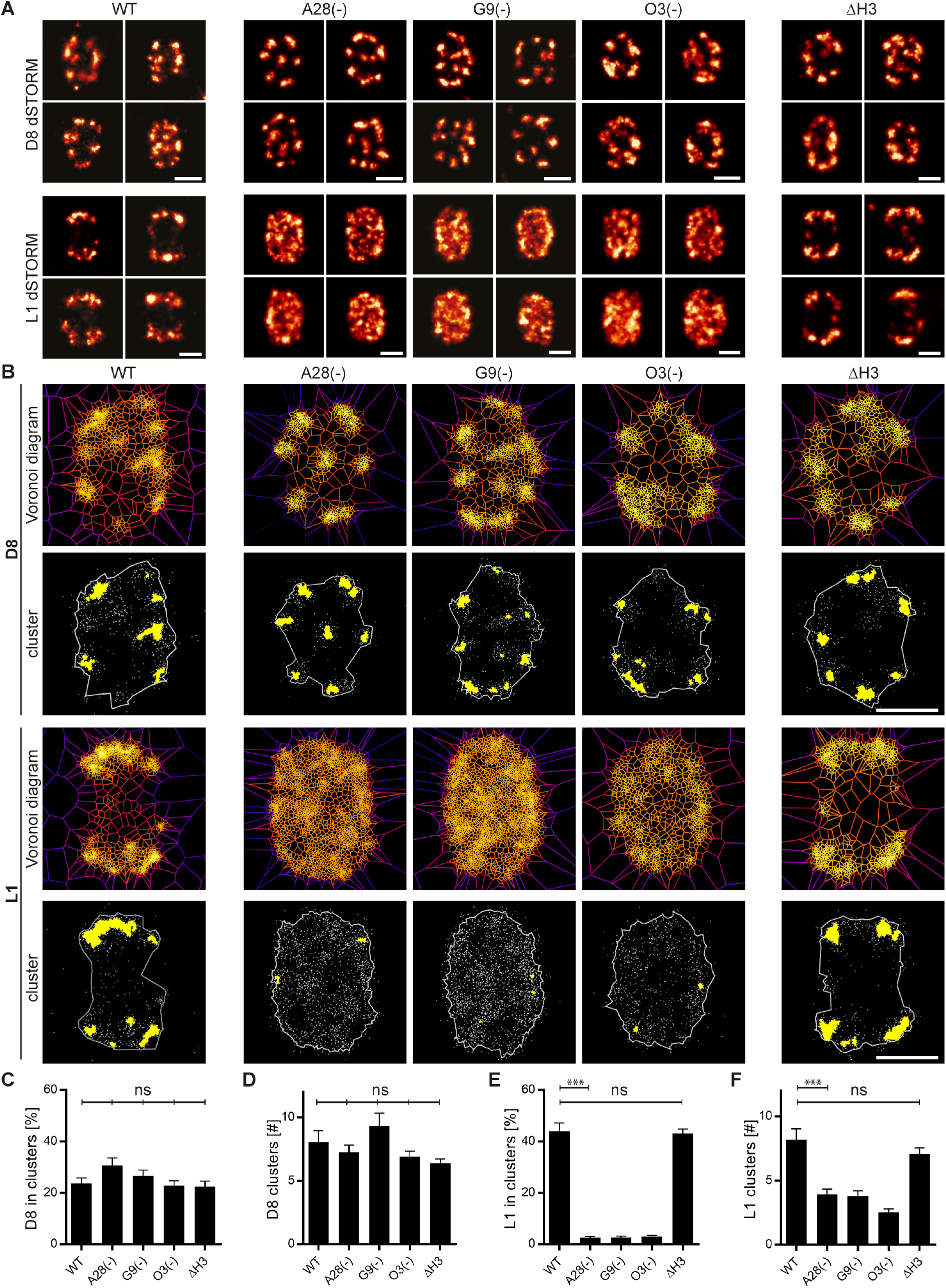
VACV binding and fusion proteins are organized into nanoscale clusters. (A) Distribution of D8 (binding protein; first row) and L1 (fusion protein; second row) on individual virions imaged by dSTORM in WT, EFC mutant [A28(-), G9(-), O3(-)], and binding protein mutant (ΔH3) MVs. (B) Voronoi diagrams and cluster identification of D8 and L1 in WT, EFC mutant, and binding mutants in (A). Voronoi tessellation was performed on individual virions with SR-Tesseler software (first row) and clusters identified with a density factor s = 3 (second row). (C) Percentage of D8 localizations within clusters on individual WT and mutant MVs. (D) Number of D8 clusters identified per virion. (E) Percentage of L1 localizations within clusters on individual WT and mutant MVs. (F) Number of L1 clusters identified per virion. Mean ± SEM, *n* = 15. Unpaired t test for significance. ***P < 0.001, ns P > 0.05. Scale bars 200nm.

To analyse D8 and L1 clustering we applied Voronoi tessellation [SR-Tesseler software (27)]. To determine a baseline for clustering, A13-EGFP (Fig. 1A) was labelled with fluorescently conjugated anti-EGFP nanobodies and imaged by dSTORM (Fig. S4D). Images were subjected to localization, segmentation and cluster identification by SR-Tesseler (Fig. S5A). The clustering threshold was set for each individual particle at 3-fold the average density (Fig. S5B). This analysis was used to quantify differences in D8 and L1 clustering between WT, EFC [A28(-), G9(-), O3(-)], and binding (ΔH3) mutant MVs (Fig. 2B). The percent of D8 localizations found in clusters, and the number of D8 clusters, showed no significant difference between WT and EFC- or binding-mutants (Fig. 2C, D). Conversely, the fraction of L1 localizations in clusters was reduced from 44% to 3%, and L1 clusters from 8 to ≤4, on WT vs. EFC mutant MVs (Fig. 2E, F, S5C, D). The total cluster area of L1 was reduced from 4,400nm^2^ /virion on WT and ΔH3 MVs, to below the imaging resolution on A28(-), G9(-) and O3(-) virions (Fig. S5E). These results show that both fusion machinery polarization and clustering are lost on EFC mutants; at the functional level this phenotype correlates with the fusion defects exhibited by these viruses (8, 24).

That A28(-), but not G9(-) or O3(-), MVs are competent for hemifusion (24), led us to hypothesize that EFC polarization and clustering play a critical role in VACV full fusion. To assess this, a viral protein that is not a component of the EFC, whose loss affects MV fusion, was needed. The VACV protein A27 has been implicated in multiple roles in the virus lifecycle including virus binding, virus fusion and the intracellular transport of virions (7, 28–31). Although A27 is a MV membrane component, it lacks a TM domain, and is tethered to the MV surface via the VACV membrane protein A17 (29). Direct evidence of the role of A27 in fusion comes from Vazquez and Esteban, who showed that A27(-) virions cannot mediate acid-induced cell-cell fusion, so-called fusion from without (32). This phenotype was confounded by the fact that the A27(-) virus has no defect in virus production (31). Confirmatory 24h-yield and fusion from without experiments with A27(+) and A27(-) MVs showed that A27(-) MVs display no defect in MV production (Fig. 3B), but are 8-fold less capable of mediating cell-cell fusion at pH 5.0 than A27(+) MVs (Fig. 3C, D).

**Fig. 3.**
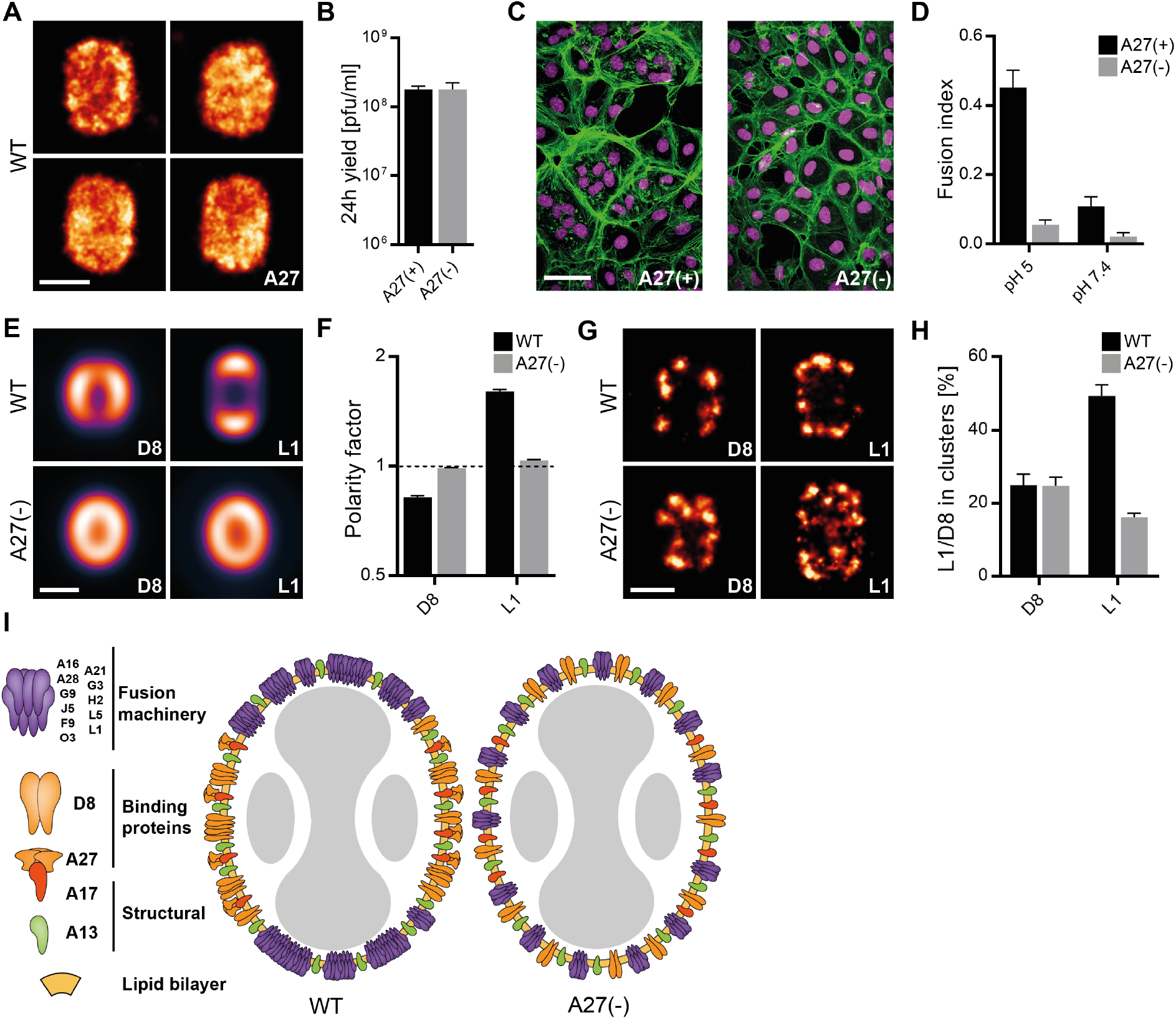
A27 regulates the protein architecture of MV membranes. (A) dSTORM images of A27 on individual MVs. (B) 24h yields of A27(+/−) virus in BSC40 cells. *n* = 3. (C) Confocal images of A27(+/−) fusion from without experiments (pH 5.0). Cell nuclei (magenta) and actin (green). Experiments were done at MOI 10. (D) Fusion index calculated for fusion from without experiments in (C). (E) VirusMapper models of L1 and D8 localization on WT and A27(-) MVs. (F) Polarity factors of models in (E). (G) dSTORM images of L1 and D8 on WT and A27(-) MVs. (H) Percent of L1 and D8 clustered localizations on WT and A27(-) MVs. *n* = 10. (I) Illustration of VACV membrane protein organization in WT vs. A27(-) MVs. Mean ± SEM. Scale bars 200nm (A,E,G) and 50 μm (C).

Considering this, we generated localization models of D8 and EFC proteins (L1, A21, A28, F9, H2 and J5) on A27(-) MVs. The distribution of D8 and EFC proteins appeared shifted on these virions (Fig. 3E, S6A). Notably, A27(-) MVs phenocopied EFC mutants, displaying a complete loss of fusion protein polarization (Fig. 3F, S6B). dSTORM showed that D8 clusters appeared redistributed, while L1 clusters were both reduced and redistributed on A27(-) MVs (Fig. 3G, H). On average, the number of L1 clusters was reduced from 8 to 6, and the total L1 cluster area reduced from 4,383nm^2^ on WT to 1,411nm^2^ on A27(-) virions (Fig. S7A, B). Conversely, no significant difference in the number of clusters or cluster area covered by D8 was detected (Fig. S7C, D). Ripley’s H-function analysis applied to L1 in WT, A27(-) and G9(-) MVs validated and confirmed these results (Fig. S7E) (33). These data indicate that A27 is required for polarized clustering of the EFC on the tips of MVs (Fig. 3I). Collectively, our results show that fusion machinery polarization requires both EFC intactness and A27. As A27 is not an EFC component, these results strongly suggest that the loss of EFC polarization in A27(-) virions underlies their inability to mediate fusion from without.

As this process relies on full fusion, as evidenced by cytoplasmic mixing experiments (24, 34). A27(+) and A27(-) MVs containing an EGFP-core were labelled with the selfquenching membrane dye R18 (34). As illustrated in Figure 4A, MVs were bound to cells in medium at 4°C/pH 7.4. This was then replaced with 37°C/pH 5.0 medium to induce fusion with the plasma membrane. This results in pH-sensitive quenching of EGFP core fluorescence and subsequent R18 dequenching due to lipid mixing during hemifusion. Upon full fusion, cores are exposed to cytosolic pH allowing for recovery of EGFP core fluorescence. Measurement of the R18 and EGFP fluorescence over time allows quantitative assessment of MV hemi- and full fusion (24, 34). Comparison of R18 dequenching rates indicated that A27(+) and A27(-) MV hemifusion was comparable (Fig. 4B). However, while core EGFP quenching occurred in both A27(+) and A27(-) MVs, only A27(+) MVs displayed significant EGFP recovery over time (Fig. 4C). This indicated that A27(-) MVs can direct hemifusion but are highly impaired in their ability to mediate fusion pore formation.

**Fig. 4.**
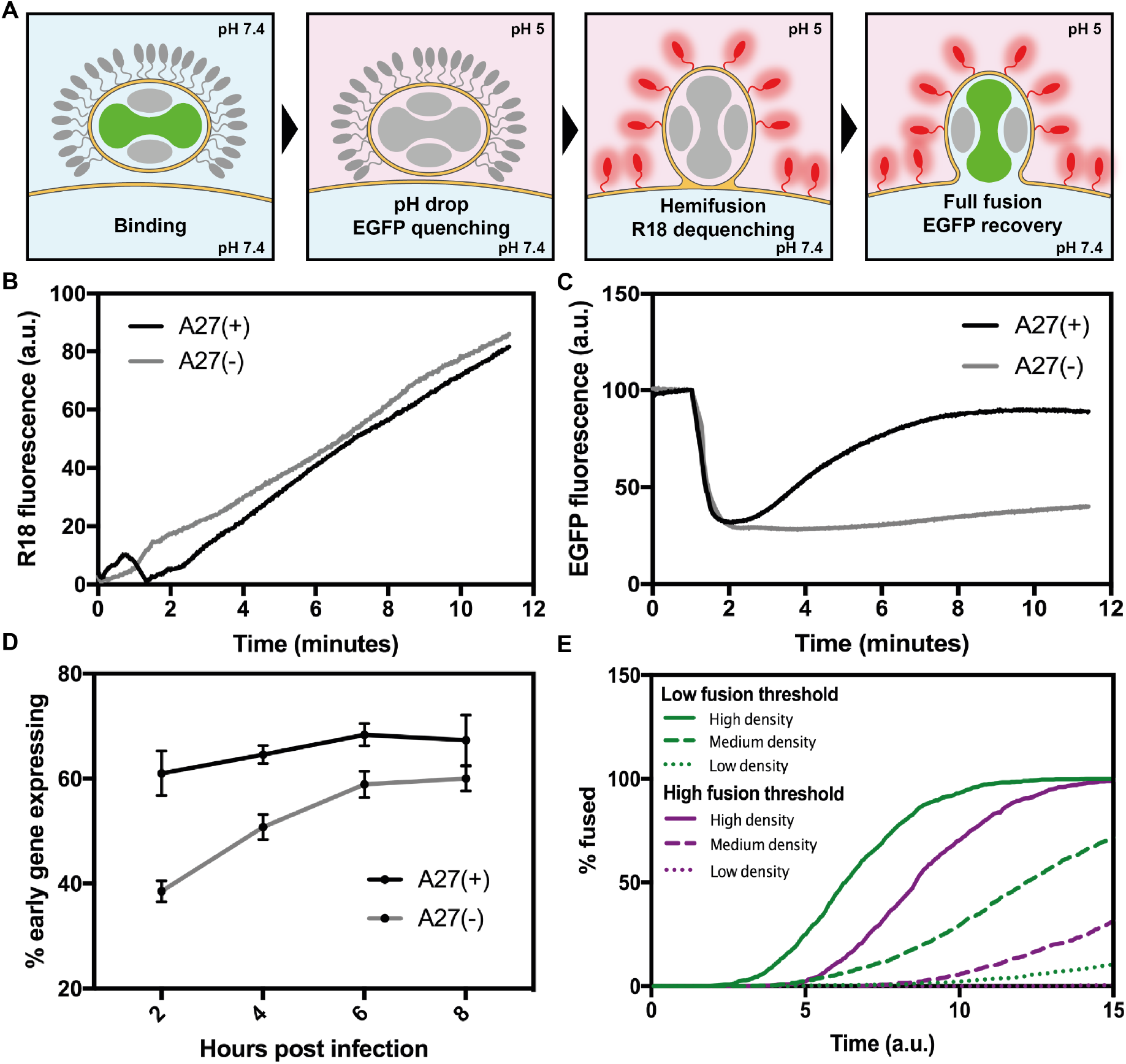
A27 regulates the protein architecture of MV membranes. (A) dSTORM images of A27 on individual MVs. (B) 24h yields of A27(+/−) virus in BSC40 cells. n=3. (C) Confocal images of A27(+/−) fusion from without experiments (pH 5.0). Cell nuclei (magenta) and actin (green). Experiments were done at MOI 10. (D) Fusion index calculated for fusion from without experiments in (C). (E) VirusMapper models of L1 and D8 localization on WT and A27(-) MVs. (F) Polarity factors of models in (E). (G) dSTORM images of L1 and D8 on WT and A27(-) MVs. (H) Percent of L1 and D8 clustered localizations on WT and A27(-) MVs. n=10. (I) Illustration of VACV membrane protein organization in WT vs. A27(-) MVs. Mean ± SEM. Scale bars 200nm (A,E,G) and 50μm (C).

We next asked how A27(-) MVs can produce equivalent numbers of infectious virus as A27(+) over 24h, despite being impaired for full fusion. Early gene expression (EGE) in cells infected with A27(+) or A27(-) MVs from 2-8 hours post infection (hpi) was compared (Fig. 4D, S8). At 2hpi, 61% of A27(+) infected cells were expressing early genes, as opposed to 38% of A27(-) infected cells. While EGE in the A27(+) infected population climbed only slightly over the 8h, in the A27(-) population EGE climbed steadily, nearly reaching the levels of A27(+) within 8h. The ability of A27(-) MVs to overcome their delayed full fusion kinetics (Fig. 4C) is consistent with the production of equivalent numbers of MVs in A27(+) and A27(-) infections over a 24h period (Fig. 3B). The experimental data indicated that clustered polarization of the fusion machinery is important for MV full fusion efficiency. To further explore the impact of EFC clustering on fusion we implemented a simple kinetic model (35). The contact area between the virus and cell was modelled to contain a variable density of fusion complexes which stochastically activate to drive fast hemifusion followed by rate-limiting full fusion. By varying the fusion complex density within the virus-cell contact area, we could assess the impact of clustering on the rate of full fusion. We tested this model at two fusion thresholds (35), low and high, which require 3 and 5 activated complexes, respectively. Simulations of 1,000 viruses indicated that the kinetics of fusion, at both fusion thresholds, are slowed when fusion complex density is reduced (Fig. 4E). Notably, at the high fusion threshold, the rate of fusion was more dependent on density. This suggests that once hemifusion occurs, full fusion is highly dependent on local fusion complex density.

## Discussion

Based on these collective findings, we propose that polarization and clustering of the poxvirus EFC is critical for efficient virus-cell fusion and MV entry. Using A27(-) MVs, which exhibit normal hemifusion but delayed full fusion kinetics, we demonstrate that EFC clustering is a major driver of full fusion efficiency, a conclusion supported by our mathematical model. To our knowledge, the only other documented example of polarized virus fusion machinery is on HIV-1 (12). In this case, a single Env cluster formed on mature virions is required for efficient CD4 engagement and entry. There, the authors proposed that Env clustering is important due to the low number of Env trimers present in the viral membrane (12). For VACV it is possible that the positioning of the fusion machinery on both virion tips serves to increase the chances of EFC-host membrane contact, successful fusion and productive infection. By revealing the nanoscale organization of the poxvirus membrane and showing the consequences of its disruption, we demonstrate that virion protein architecture is critical to virus function. We suggest that the organization of the VACV membrane into functionally distinct domains has evolved as a mechanism to maximise virion binding and fusion efficiency for productive infection.

## ACKNOWLEDGEMENTS

We would like to thank Bernard Moss and Mariano Esteban for kindly providing mutant viruses used in this study. This work was funded by MRC Programme Grant (MC-UU12218/7) (J.M.), the European Research Council (649101-UbiProPox) (J.M.), the UK Medical Research Council (MR/K015826/1) (J.M., R.H.) Biotechnology and Biological Sciences Research Council (BB/M022374/1; BB/P027431/1; BB/R000697/1) (R.H.), and the Wellcome Trust (203276/Z/16/Z) (R.H.). R.G. is funded by the Engineering and Physical Sciences Research Council (EP/M506448/1), D.A. is presently a Marie Curie fellow (Marie Sklodowska-Curie 750673). C.B is funded by the Medical Research Council LMCB PhD program.

## AUTHOR CONTRIBUTIONS

R.G., D.A., R.H. and J.M. conceived the project, designed the experiments and wrote the manuscript. R.G., D.A., and C.B performed the experiments. The Cohen (G.H.C.) and Eisenberg members of the poxvirus research group produced and purified all VACV EFC antibodies. R.G., D.A., analysed the data. R.G., D.A., R.H. and J.M. discussed the results and implications of the findings. All authors discussed and provided comments on the manuscript.

## Methods

### Cell lines and viruses

African green monkey kidney (BSC-40) cells and human HeLa cells were cultivated in Dulbecco’s modified Eagle’s medium (DMEM) supplemented with 10% heat-inactivated fetal bovine serum (FBS), 2mM Glutamax, 100units/mL penicillin and 100(μg/mL streptomycin, 100(μM nonessential amino acids and 1mM sodium pyruvate. Routine mycoplasma tests of the cell culture medium were negative. Recombinant VACV were based on VACV strain Western Reserve (WR). WR mCherry-A4 and WR L4-mChen-y/EGFP-F17 were described previously as WR mCherry-A5 (1) and WR EGFP-F17 VP8-mCherry (2). ΔH3, A28(+/−), G9(+/−) and O3(+/−) were described previously as vH3Δ (3), vA28-HAi (4), vG9i (5) and v03-HAi (6) respectively. WR A27(+/−) was described previously as VVIndA27L (7). WR A4-mCherry/A13-EGFP, WRA27(+/−) EGFP-A4 and WR A27(+/−) E/L EGFP were constructed as previously described (2). For WR A4-mCheny/A13-EGFP, A13 was replaced with A13-EGFP in its endogenous locus within WR A4-mCherry virus. For WR A27(+/−) EGFP-A4, A4 was replaced with EGFP-A4 in its endogenous locus within WR A27(+/−). For WR A27(+/−) E/L EGFP, EGFP under the control of an early/late viral promoter was inserted between the H2-H3 loci of the WR A27(+/−) virus. Briefly, BSC-40 cells were infected the parental viruses and subsequently transfected with linearized plasmid containing the region of genome to be replaced, flanked 5’ and 3’ by 300bp of genomic sequence for targeted homologous recombination. Recombinant viruses were selected by fluorescence through 4 rounds of plaque purification. All viruses were produced in BSC-40 cells, and MVs purified from cytoplasmic lysates by banded on sucrose gradients, as previously described (8). For WR A27(+/−) and its derivatives, virus stocks were generated in the presence or absence of 2mM Isopropyl *β*-D-thiogalactoside (IPTG, Sigma) to obtain A27(+) or A27(-) MVs. WR A28(+), WR G9(+) and WR O3(+) stocks were produced with 100(*μ*M, 50(*μ*M and 20(*μ*M IPTG, respectively.

### Antibodies

Anti-L1 mouse monoclonal antibody (clone 7D11) was purified from a hybridoma cell line kindly provided by Bernard Moss (National Institutes of Health, Bethesda, MD) with permission of Alan Schmaljohn (University of Maryland, Baltimore, MD). Anti-D8 rabbit polyclonal antibody was made by immunizing a rabbit with purified D8 protein and adjuvant. Anti-A17 rabbit polyclonal antibody was a kind gift from Jacomine Krijnse-Locker. Antibodies against viral proteins A21 (R206), A28 (R204), F9 (R192), H2 (R202), J5 (R264), were produced by GHC using purified recombinant baculovirus-expressed proteins as previously described (9).

### Structured illumination imaging

High performance coverslips (18 × 18mm 1.5H, Zeiss) were washed as previously described (10). Purified virus was diluted in 1mM Tris pH 9, bound to the ultra-clean coverslips for 30 min and fixed with 4% Formaldehyde. Samples were washed 3 times with PBS prior to mounting for imaging. To visualize membrane proteins, virus was blocked after fixation using 5% bovine serum albumen (BSA, Sigma) in PBS for 30 min, incubated in 1% BSA in PBS with primary antibody, followed by Alexa Fluor 488-conjugated goat anti-mouse or anti-rabbit secondary antibody for 1 hour each. Samples were washed 3 times with PBS after each staining step. The coverslips were mounted in Vecta Shield (Vector Labs) and sealed with nail polish. SIM imaging was performed using Plan-Apochromat 63x /1.4 oil DIC M27 objective, in an Elyra PS.1 microscope (Zeiss). Images were acquired using 5 phase shifts and 3 grid rotations, with the 561nm (32μm grating period) and the 488nm (32μm grating period) lasers, and filter set 3 (1850–553, Zeiss). 2D images were acquired using a sCM0S camera and processed using the ZEN software (2012, version 11.0.3.190, Zeiss). For channel alignment, TetraSpeck beads (ThermoFisher) were mounted on a slide, imaged using the same image acquisition settings and used for the alignment of the different channels.

### Single-particle analysis

Individual viral particles were extracted from the SIM images. Seed images were generated with VirusMapper as described previously (10). VirusMapper models were then created by registration of the entire set of particles according to cross-correlation with the seeds and calculation of a weighted average of a subset of particles. These values of (n) are shown in Figs. S3 and S6.

### Polarity factor

Polarity factors were calculated directly from the sets of particles used to generate the models (Fig. S2). Following the cross-validation method of Szymborska et al. (11), particles were randomly divided into subsets of 50 particles and separately averaged. Radial profiles were generated from these images by transforming from x - y coordinates to r - *θ*. The radial profiles were divided into four regions according to a parameter *ϕ* and the mean intensity within the viral membrane in these regions was evaluated. The four regions were defined in *θ* by:

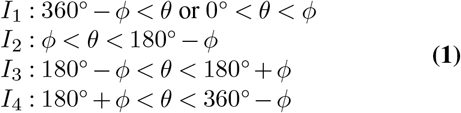

where 0 *<9 <* 360°. The images are therefore divided by two lines through the centre of the image that intersect at angle *ϕ* and divide the image symmetrically about the horizontal and vertical axes. The polarity factor was then calculated as

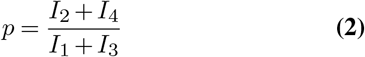

A value for *ϕ* was used that resulted in a mean polarity factor of 1 for the A4 core protein (48.5°). Values quoted in the text were calculated by averaging the mean polarity factors for the sets of binding and fusion proteins.

### Labelling of primary antibodies

Mouse monoclonal antibody (IgG) against L1 from hybridoma supernatant and rabbit polyclonal antibody against D8 from whole serum were purified with a Nab Protein A/G Spin Kit (Thermo Scientific) according to the manufacturer’s instructions. Antibodies were buffer exchanged into 0.1M NaHCO3 pH 8.2 and concentrated to >1mg/ml concentration with Amicon Ultra centrifugal filters 10 kDa MWCO (Merck). Antibodies were custom-labelled with a 50-fold molar excess with AlexaFluor647-NHS (Invitrogen) for 1h at RT. The reaction was quenched by addition of 2 *μl* 1M Tris pH 9. Unreacted dye was removed by three passes through Zeba-Spin columns 3.5K MWCO (Thermo Scientific).

### dSTORM imaging

High performance coverslips (18mm, 1.5H, Zeiss) were washed in ultrapure ethanol (Sigma) and deionized water to clean them and make their surface hydrophobic. Purified virus was diluted in 20*μ*il 1m MTris pH 9, placed in the centre of the clean coverslips for 30min and bound virus fixed with 4% Formaldehyde. Virus was blocked after fixation using 5% BSA (Sigma), 1% FCS, 0.2-1% TritonX-100 in PBS for 30 min. Virus was immunostained in 5% BSA in PBS with primary antibodies at 4 °C overnight and Alexa Fluor 647-conjugated secondary antibodies (Invitrogen) for 1-2 hours at RT. Samples were washed 3 times with PBS after each staining step. Optionally, a post-fixation step with 4% PFA was included. Autofluorescence was quenched by brief incubation in 0.25% (w/v) NH4Cl in PBS. Coverslips were mounted on a Secure-Seal incubation chamber (EMS) in BME-buffer [1% (v/v) β-mercaptoethanol (Sigma), 150 mMTris, 1% glucose, 1% glycerol, 10mM NaCl, pH 8] or in MEA-buffer [50mM cysteamine (Sigma), 3% (v/v) OxyFluor (Oxyrase Inc), 20% sodium lactate (Sigma), PBS pH 8 (12)]. dSTORM imaging was performed on an Elyra PS.1 inverted microscope (Zeiss) using an alpha Plan-Apochromat 100x/1.46 NA oil DIC M27 objective with a 1.6x tube lens and an iXon 897 EMCCD camera (Andor). Images were acquired at 25-30ms exposure time with 642nm excitation at 100% laser power and a 655nm LP filter. Fluorophore activation was dynamically controlled with a 405nm laser at 0–2% laser power. Images were processed in Fiji (13) using ThunderSTORM (14). Localizations were fitted with a maximum-likelihood estimator, lateral drift corrected by cross-correlation, localizations < 20nm apart within ≤ 1 frames merged, and images rendered using a Gaussian profile with the NanoJ-Orange LUT (NanoJ). Lateral resolution was 25nm, determined by FRC.

### SR-Tesseler analysis

Cluster analysis was performed with SR-Tesseler (15). Localization tables from ThunderSTORM were imported and Voronoi diagrams created. Individual virions were selected as regions of interest and segmented as single objects with a density factor *6* of 0.1 – 0.5. Within individual objects, clusters were identified with *6* = 3 which yielded <2% clustering in the non-clustered reference probe A13-EGFP (Fig. S5). Statistical analysis was performed in GraphPad Prism (Prism Software). Significance (unpaired t-test): * P < 0.05; ** P <0.01; *** P < 0.001.

### Ripley’s H-function analysis

Additional cluster analysis was carried out as in (16). 10 representative virus particles were selected from dSTORM images of L1 on WT, A27(-), G9(-) VACV (Fig. S7E). Localizations within the selected ROIs were separately used to calculate the Ripley’s H-function as a function of increasing radius H(r) according to:

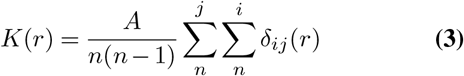

where *A* is the approximate area of the particle, *n* is the number of localizations and

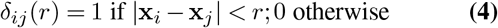

where *x_i_* is the spatial location of localization i. Then

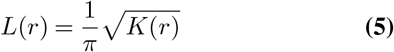

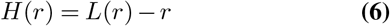

so that the expected value of *H(r)* is 0 for a random uniform distribution, and values of *H(r)* above 0 indicate clustering at a range of approximately r. Custom Python scripts were used to calculate the H-function from dSTORM localization tables. The mean of *H(r)* and the 95% confidence interval were evaluated for each case.

### 24-hour yields

Confluent BSC-40 cells were infected with A27(+/−) virus in DMEM at MOI 1. After 60 minutes the inoculum was removed and DMEM with serum and 2mM IPTG was added for 24 hours. Cells were harvested, resuspended in 100*μ*il 1mM Tris pH 9 and freeze-thawed 3 times in liquid nitrogen. The virus concentration was then measured by plaque assay.

### Fusion from without

WRA27(+/−) was bound to HeLa cells grown on coverslips on ice at MOI 10 for 1 hour. Cells were washed and incubated at 37°C with 20mM MES pH 5 or 7.4 for 5 mins, then washed and incubated in full medium for 2 hours. Cells were fixed with 4% formaldehyde in PBS for 20 mins, blocked and permeabilized using 0.1% Triton X-100 and 5% BSA in PBS, stained with Alexa Fluor 488 phalloidin and Hoechst 33258 (both Invitrogen) for in PBS for 1 hour. Samples were then mounted on slides with ProLong Gold Antifade Mountant (Thermo Fisher). Samples were imaged on a Leica TCS SP8 STED 3x microscope in confocal mode running LAS X (Version 2.01) acquisition software. Alexa Fluor 488 fluorescence was excited using the 488 nm line and Hoechst was excited with the 405nm LED. Z-stacks were acquired at a scan speed of 400 Hz in bidirectional scan mode using a Hybrid Detector (HyD, Standard mode) and a Acousto-Optical Beam Splitter for filtering and time-gating of 0.5-8ns. The fusion index was calculated from maximum intensity projections of the stacks as described previously (17).

### Bulk fusion measurements

MVs were labelled by incubating with 22.5*μ*M R18 (ThermoFisher) in 1mM Tris pH 9 at room temperature for 2 hours. Labelled viruses were pelleted by centrifugation at 16,000g for 10 min at 4°C and resuspended in 1mM Tris pH 9 twice to remove excess R18. Labelled viruses were bound to 70000 HeLa cells at MOI 30 in DMEM on ice for 1 hour. Cells were sedimented by centrifugation at 300g for 5 mins at 4°C then resuspended in 100*μ*l PBS. The cell suspension with bound virions was added to 630*μ*l prewarmed PBS in a quartz cuvetre. After 2 min, the pH in the cuvetre was lowered by addition of 100*μ*l 100mM MES resulting in a pH of 5.0. After the acquisition all R18 was dequenched by the addition of 83*μ*l 10% Triton X-100 in PBS. R18 fluorescence was normalized to the signal intensity after Triton X-100 addition. R18 fluorescence was measured using a Horiba FluoroMax 4 (Horiba Jobin Yvon) spectrofluorometer with an excitation wavelength of 560 ± 5 nm and an emission wavelength of 590 ± 5 nm. To measure EGFP fluorescence, unlabelled viruses were bound to cells and the pH was lowered as above, and fluorescence was recorded using an excitation wavelength of 488 ± 5 nm and an emission wavelength of 509 ± 10 nm.

### Structured illumination imaging

Flow cytometry HeLa cells were infected with WR A27(+/−) E/L EGFP at MOI 4 in DMEM and full medium containing 10µM AraC (Sigma) was added after 1 hour. At the indicated times cells were washed with PBS, trypsinized and resuspended in PBS and fixed with 4% formaldehyde in PBS. Cells were then sedimented by centrifugation at 300g for 5 mins and resuspended in PBS with 2% FBS and 5mM EDTA. Flow cytometry was performed with a Guava EasyCyte HT flow cytometer, recording the EGFP fluorescence with the 488nm laser. Analysis of the flow cytometry data was performed with the FlowJo software package.

### Fusion kinetics mathematical model

We modelled the process of fusion undergone by a single fusion complex as a fast irreversible hemifusion step followed by a rate-limiting, irreversible full fusion step.

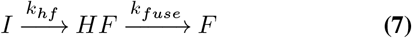

Following similar work on influenza (18), for each virus we required that *T_f_* (fusion threshold) fusion complexes transition to state F before the virus is considered fused. We simulated the contact area with the cell as containing an initial density po (high density), 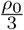 (medium density) and 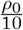 (low density) of fusion complexes. We then modelled the effects of fusion complex density on fusion kinetics at two different values of Tf: low (*T_f_* =3) and high (*T_f_* = 5). The true value of *T_f_* is unknown for vaccinia but for influenza is thought to be in this range.

---

**Fig. S1.**
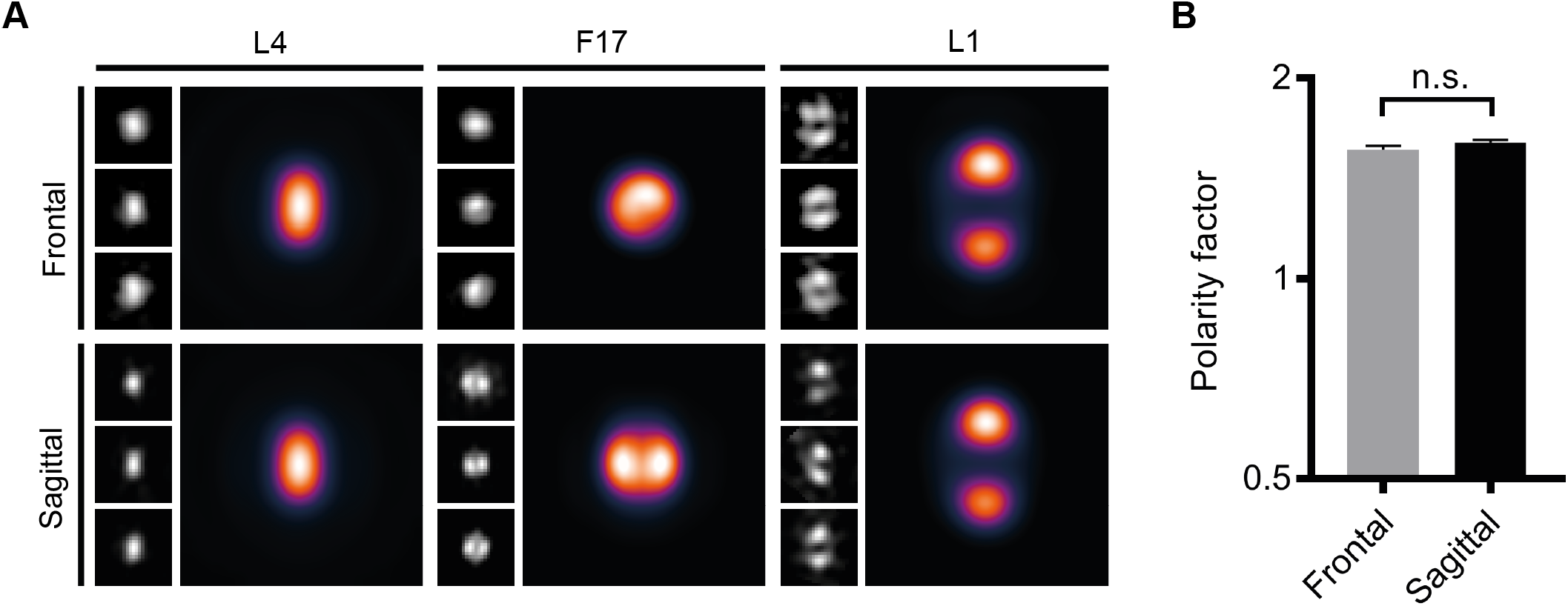
Fusion protein polarisation is independent of lateral orientation. (A) Viruses with core protein L4 tagged with mCherry and lateral body protein F17 tagged with EGFP (WR L4-mCherry F17-EGFP) were labelled for fusion protein L1 and imaged with SIM. According to the appearance of the lateral bodies, particles were assigned to either frontal or sagittal orientation and averaged using VirusMapper. (B) Polarity factors were calculated for the frontal and sagittal case. Shown is the mean ± SEM of the polarity factor values for separately averaged random subsets of 50 particles with unpaired t-test. ns P > 0.05.

**Fig. S2.**
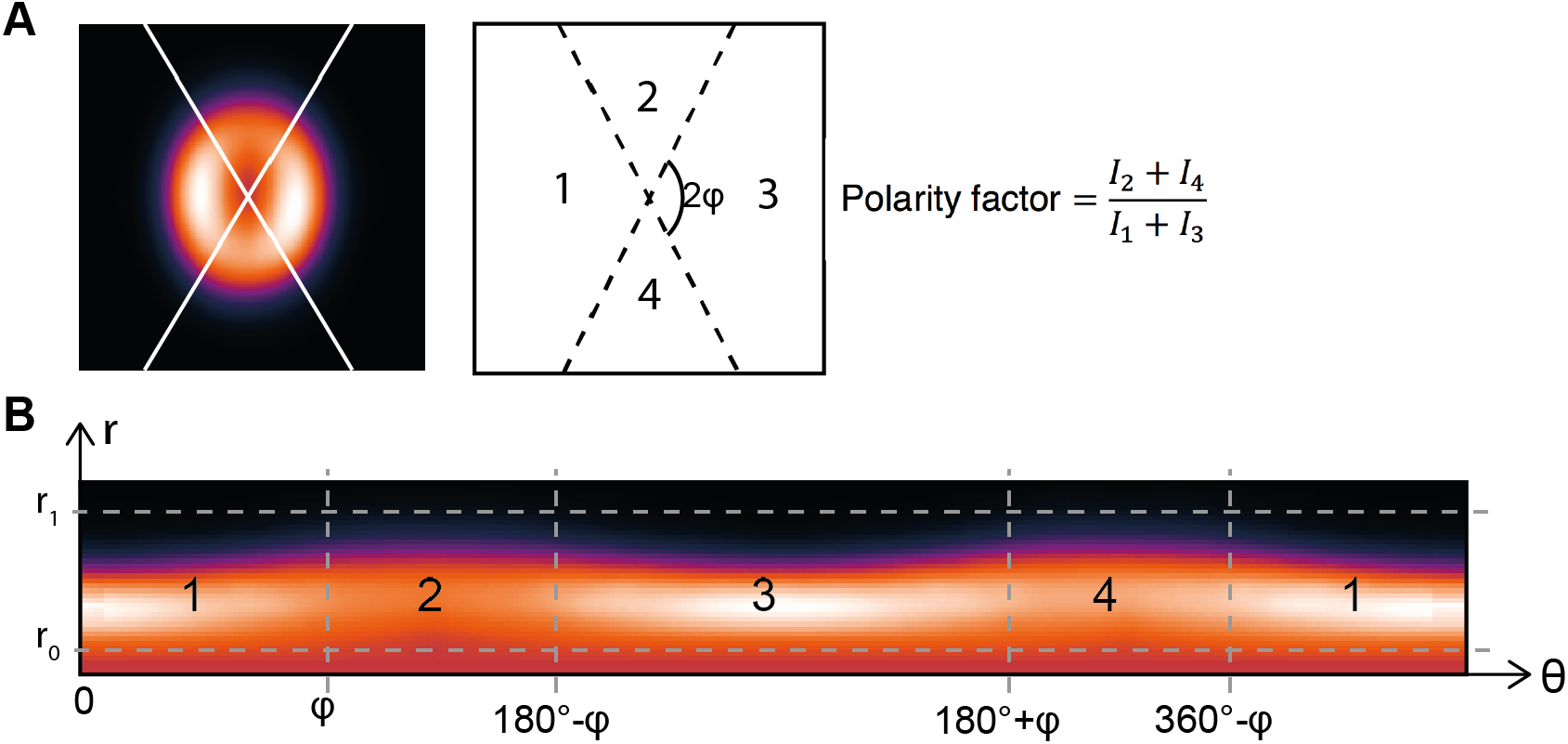
Quantification of membrane protein polarisation. (A) Images of viruses stained for proteins of interest were split into four sections, and the polarity factor was calculated as the ratio of the sum of the intensities in the polar sections to the sum of the intensities in the lateral sections. (B) This was done by first calculating a radial profile of the image and then evaluating the intensities in the radial profile. The radius was restricted to the membrane region for membrane proteins and the section boundaries, defined by a single angle *φ*, were set so that core protein A4 displayed a polarity factor of 1.

**Fig. S3.**
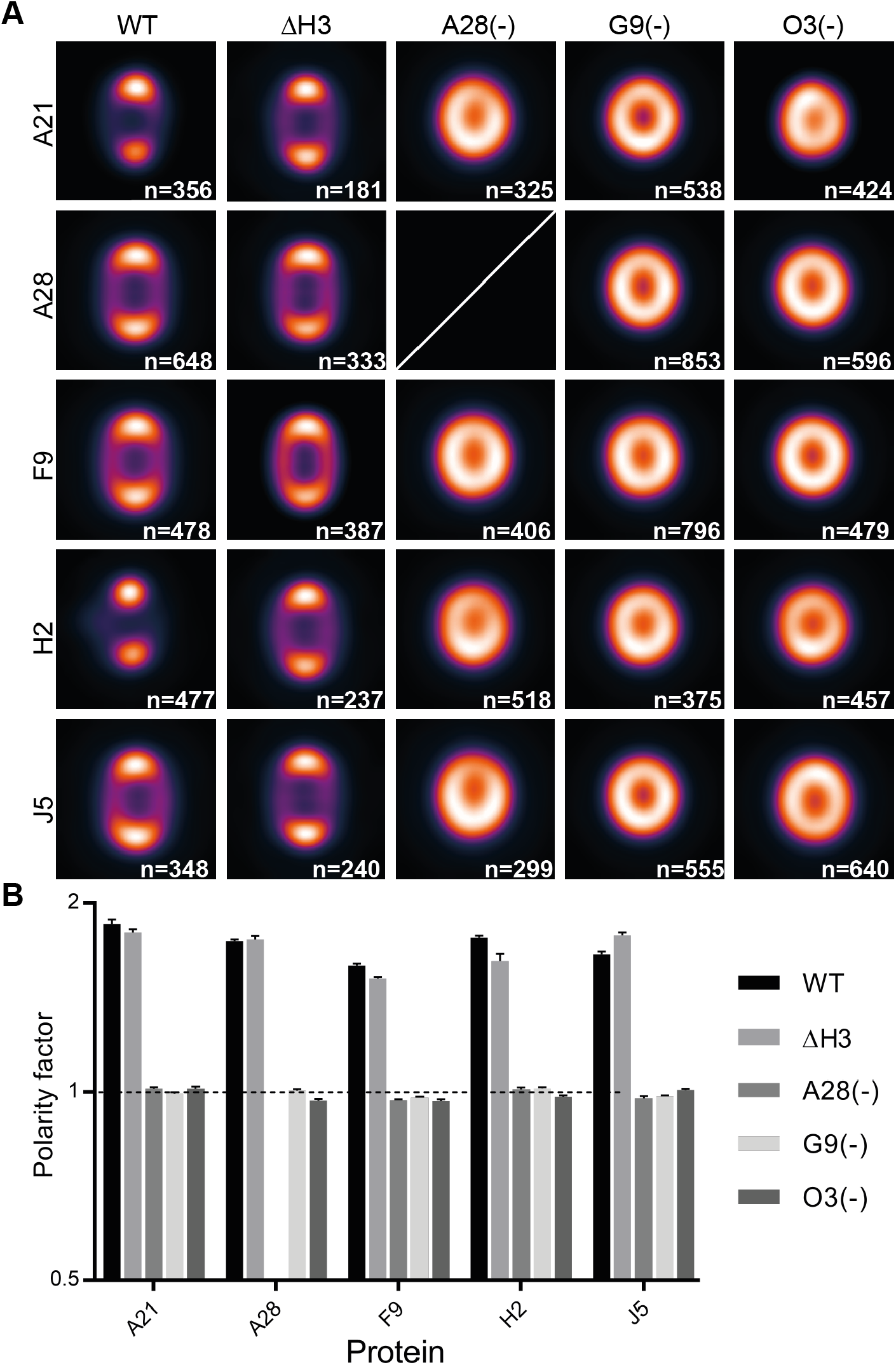
VirusMapper modelling of fusion machinery proteins In EFC mutant viruses. (A) SIM-based VirusMapper models of the localizations of 5 fusion machinery proteins in WT, 3 EFC- and 1 binding- mutant virus. The number of particles averaged for each model is displayed. No model was generated for A28 in the A28(-) virus as the protein is not present. (B) Polarity factors for the models in (A). Mean ± SEM for separately averaged subsets of 50 particles.

**Fig. S4.**
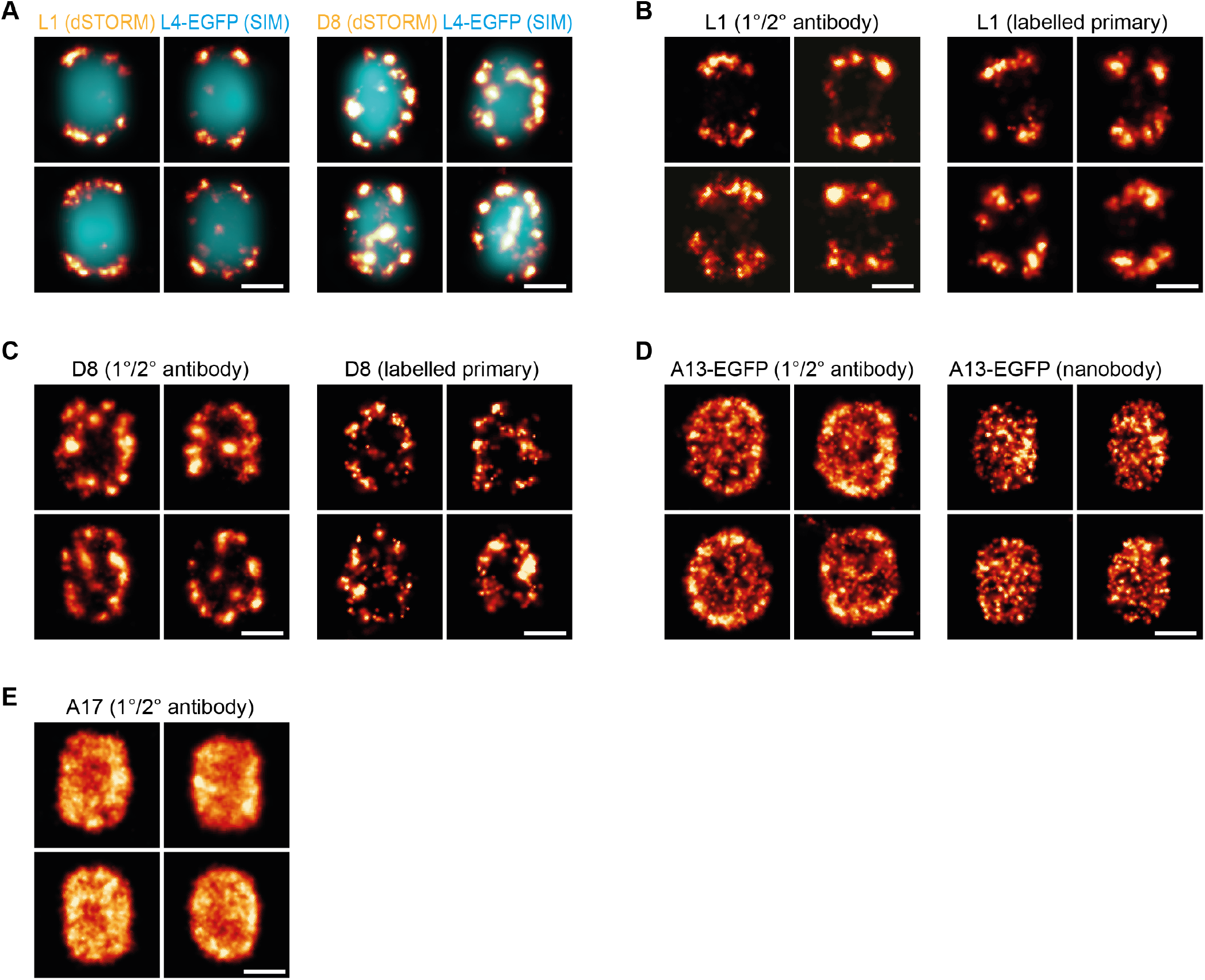
Super-resolution imaging of viral proteins. (A) Correlative dSTORM and SIM imaging of viral proteins. VACV core protein L4-EGFP was imaged by SIM (cyan) to determine the orientation of the viral particle. EFC marker L1 or binding protein D8 were then imaged by dSTORM (LUT: NanoJ-Orange) and images overlaid in imageJ using NanoJ image registration. (B) dSTORM images of WT vaccinia virus immunolabelled with L1 primary antibody and secondary anti-mouse antibody conjugated to AF647, or with L1 primary antibody directly conjugated to AF647. (C) dSTORM images of WT vaccinia virus immunolabelled with D8 primary antibody and secondary anti-rabbit antibody conjugated to AF647, or with D8 primary antibody purified from serum and directly conjugated to AF647. (D) Localization of A13-EGFP using polyclonal anti-GFP antibody and secondary anti-rabbit AF647 or with the single-binder anti-GFP nanobody conjugated directly to AF647. Notably, the virus appears bigger when labelled with antibodies compared to nanobodies, due to the different label sizes. (E) dSTORM images of VACV protein A17. Scale bars = 200nm.

**Fig. S5.**
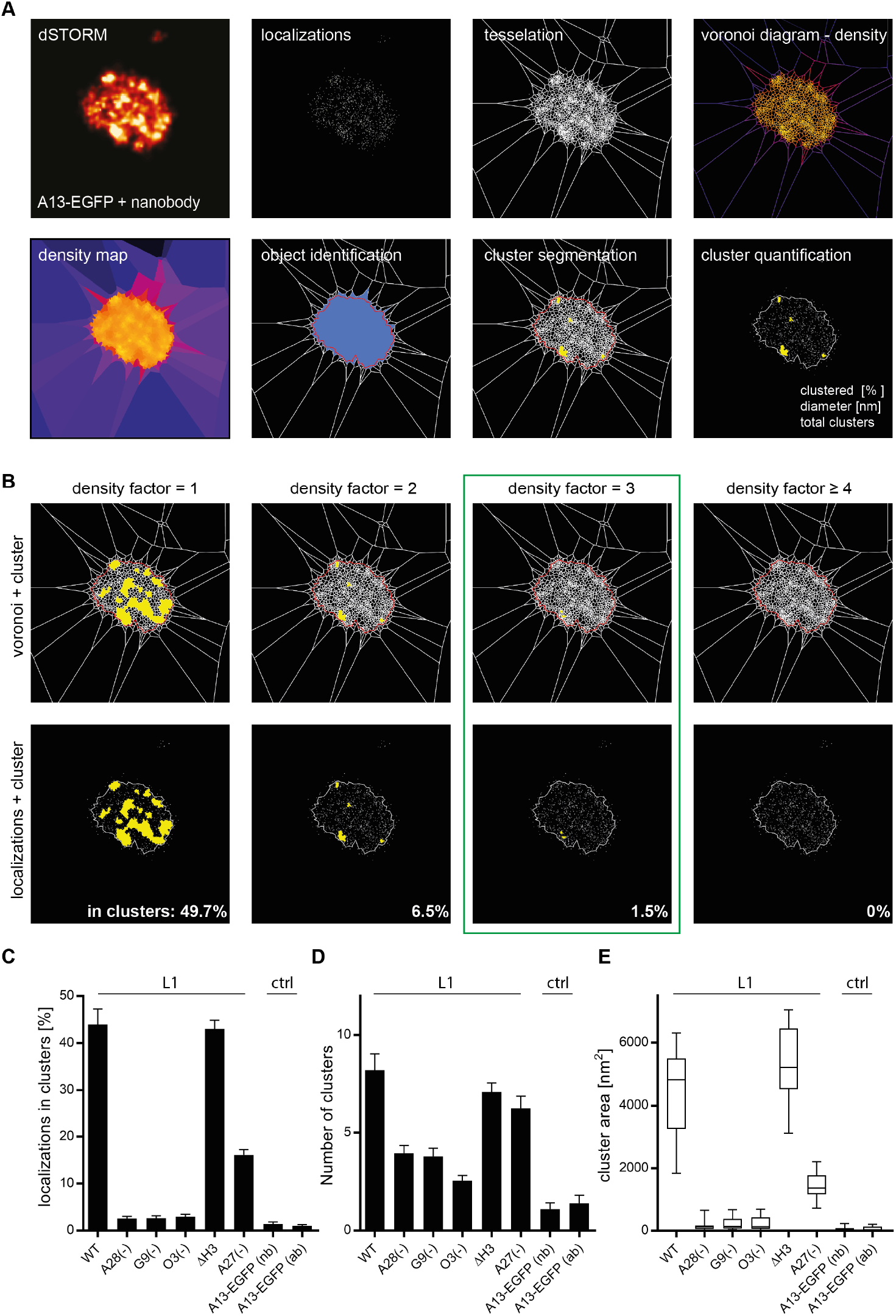
Cluster segmentation and analysis with SR-Tesseler software. (A) Images of A13-EGFP labelled with single-binder anti-GFP nanobodies acquired by dSTORM. Localization tables were imported into SR-Tesseler software and a voronoi diagram created via tessellation. The density map with tiles color-coded for density is an alternative illustration of localization data comparable to the reconstruction via gaussian profiles in the first panel. Individual virions were cropped as regions of interest and identified as objects in SR-Tesseler. Clusters were segmented in individual objects based on a density factor δ. The percentage of localizations attributed to clusters, cluster diameter and total number of clusters per object were quantified. (B) Cluster identification was performed with density factors δ ranging from 1 (equal to the average density of the object) to 4 (four-fold higher density than average). At δ = 1 approx. 50% of localizations were attributed to clusters, as expected. δ = 3 was selected as the threshold for all data analysis with < 2% clustering of the per definition unclustered A13-EGFP Higher thresholds (δ ≥ 4) caused underrepresentation of clustering. (C) Localizations in clusters for L1 in WT and mutant [A28(-), G9(-), O3(-), ΔH3, A27(-)] MVs and for A13-EGFP labelled with nanobodies (nb) or antibodies (ab). MVs lacking an intact EFC show residual clustering similar to the A13-EGFP control. (D) Average number of clusters for L1 in WT and mutant [A28(-), G9(-), O3(-), ΔH3, A27(-)] MVs and for A13TM-EGFP labelled with nanobodies (nb) or antibodies (ab). (E) Cluster area for L1 in WT and mutant [A28(-), G9(-), O3(-), ΔH3, A27(-)] MVs. Cluster area is negligible in EFC deficient MVs as in the unclustered control A13-EGFP Error bars represent SEM (C,D) and 95th percentile (E).

**Fig. S6.**
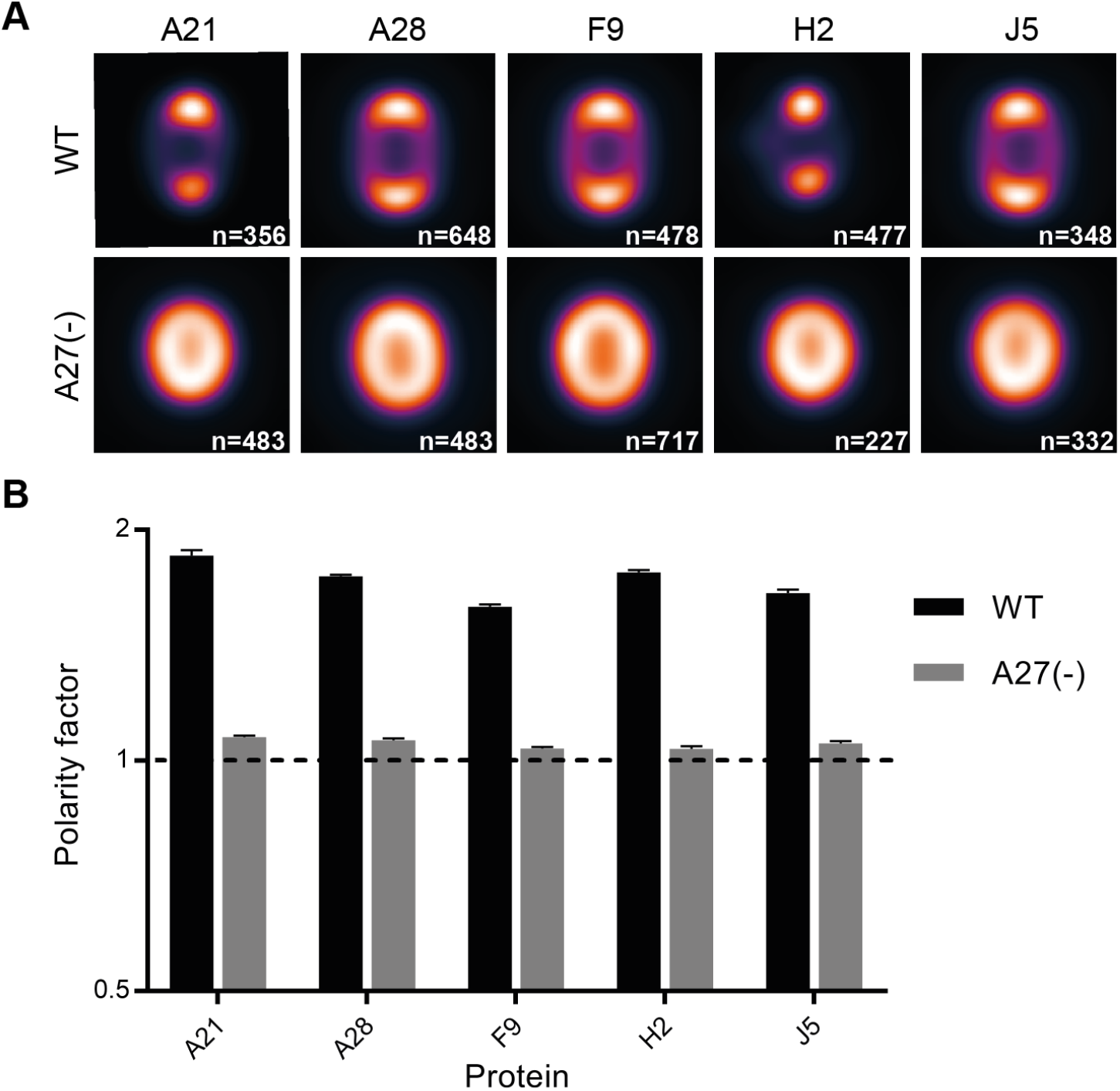
VirusMapper modelling of fusion machinery proteins in A27(-) virus. (A) SIM-based VirusMapper models of the localizations of 5 fusion machinery proteins in A27(-) virus compared to WT. The number of particles averaged for each model is displayed. (B) Polarity factors of models in (A). Mean ± SEM for separately averaged subsets of 50 particles.

**Fig. S7.**
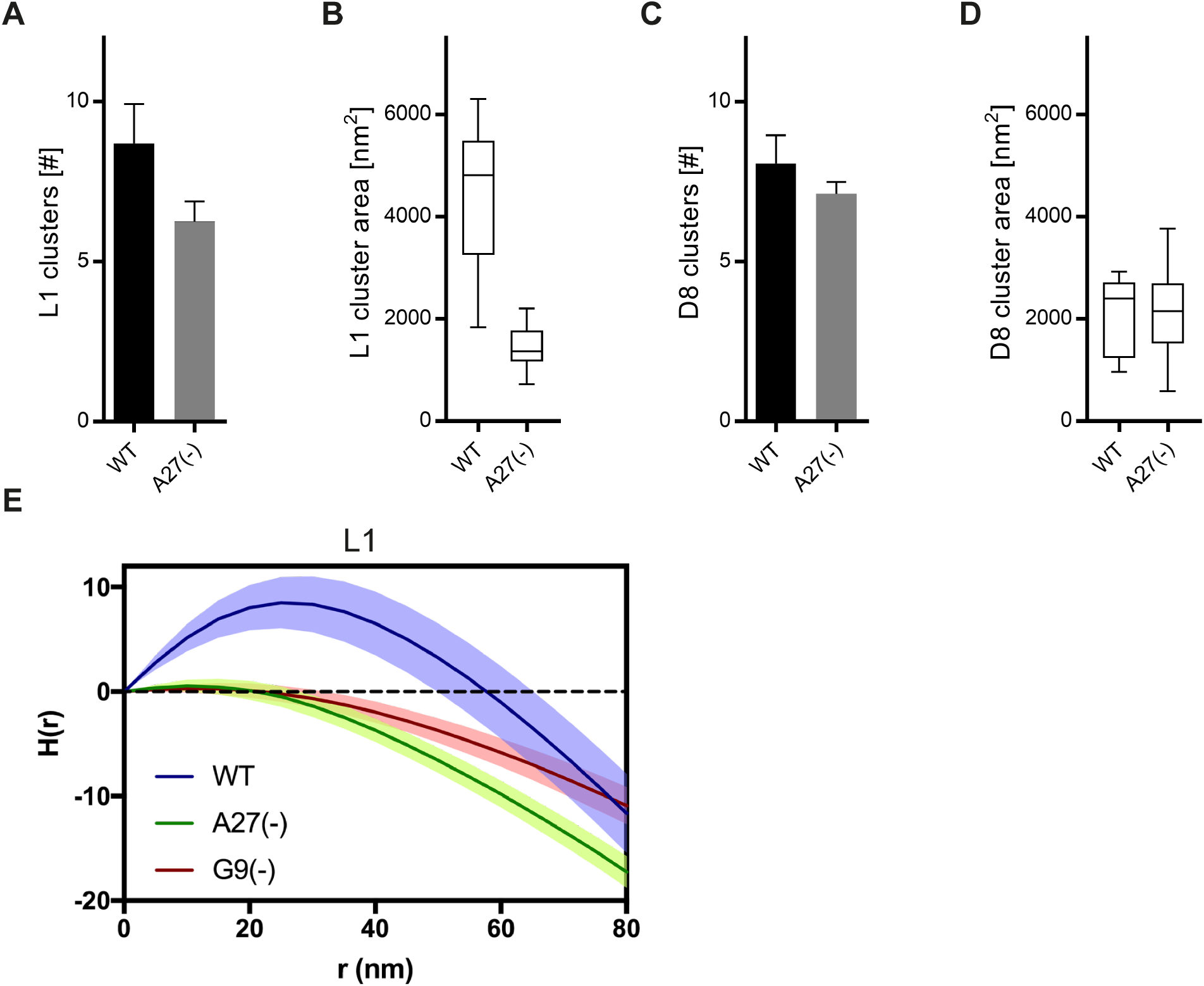
Quantification of L1 and D8 clustering in A27(-) MVs via SR-Tesseler and Ripley’s H-function analysis. (A) Cluster number for L1 in WT and A27(-) MVs, n=15. (B) Cluster area for L1 in WT and A27(-) MVs, ***n*** = 15. (C) Cluster number for D8 in WT and A27(-) MVs, ***n*** = 15. (D) Cluster area for D8 in WT and A27(-) MVs, n=15. (E) Ripley’s H-function was calculated for individual virus particles from dSTORM localization lists. A value greater than 0 indicates clustering and less than 0 indicates dispersion. Clustering of L1 is seen in WT but not G9(-) or A27(-) viruses. Mean and 95% confidence interval, n=10. Error bars SEM (A,C) and 95th percentile (C,E).

**Fig. S8.**
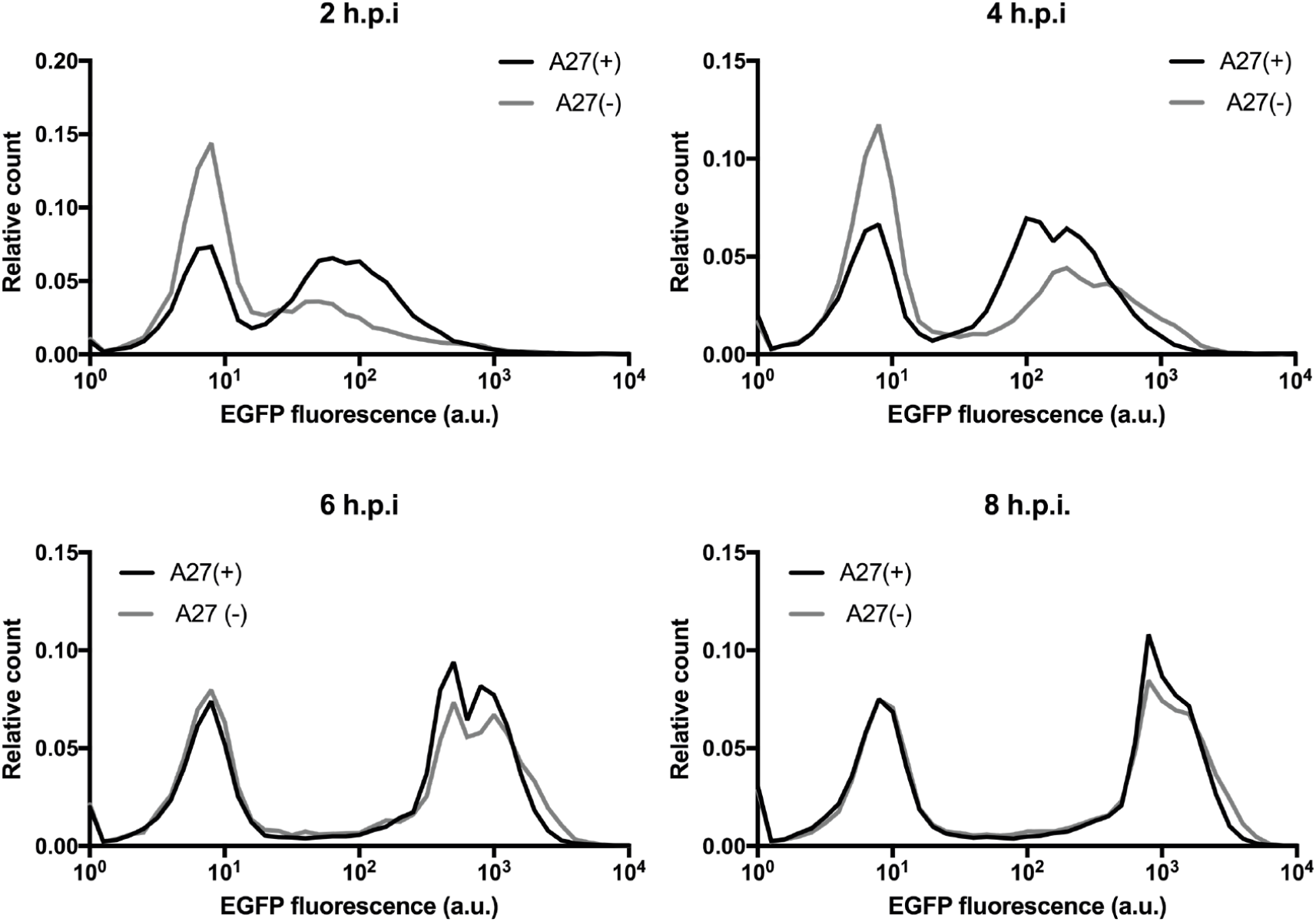
VACV early gene expression kinetics are delayed in the absence of A27. Flow cytometry histograms of the distribution of EGFP fluorescence intensity as a proxy for early gene expression over time for A27(+) and A27(-) virus.

